# Landmark-Based Turn-by-Turn Instructions Enhance Incidental Spatial Knowledge Acquisition

**DOI:** 10.1101/2020.11.30.403428

**Authors:** Anna Wunderlich, Sabine Grieger, Klaus Gramann

**Affiliations:** Technische Universität Berlin, FG Biopsychologie und Neuroergonomie, Berlin 10623, Germany; School of Computer Science, University of Technology Sydney, Sydney 2007, Australia; Center for Advanced Neurological Engineering, University of California, San Diego, CA, USA

**Keywords:** navigation assistance, spatial knowledge acquisition, virtual driving, long-term, landmarks

## Abstract

The augmentation of landmarks in auditory navigation instructions had been shown to improve incidental spatial knowledge acquisition during assisted navigation. Here, two driving simulator experiments are reported that replicated this effect even when adding a three-week delay between navigation and spatial tasks and varying the degree of detail in the provided landmark information. Performance in free- and cued-recall of landmarks and driving the route again without assistance demonstrated increased landmark and route knowledge when navigating with landmark-based compared to standard instructions. The results emphasize that small changes to existing navigation systems can foster spatial knowledge acquisition during every-day navigation.

## Introduction

Daily spatial orienting tasks like wayfinding are increasingly supported by navigation aids that provide visual as well as auditory guidance. The use of automated navigation assistance systems requires dividing attention between the locomotion tasks and the assistance system (Fenech et al., 2010; Gardony et al., 2013, 2015). To reduce the attentional demand, users increasingly rely on the navigation assistant (Baus et al., 2001; Klippel et al., 2010; Parush et al., 2007) simply following turn-by-turn instructions without processing further information available to them (Mosier et al., 1996; Parasuraman, 2000). As a consequence, spatial information from the surrounding environment is less processed and the ability to extract navigation relevant information diminishes (Aporta & Higgs, 2005; Fenech et al., 2010).

One promising approach to counter this effect of assisted navigation is the reference to salient objects, so called landmarks (Evans et al., 1982), in the navigation instructions (Goodman et al., 2005; Li et al., 2014; Löwen et al., 2019). Auditory compared to visual augmentation methods have the advantage that they do not distract the user from monitoring the ongoing traffic and control of the vehicle. Auditory augmentation of landmarks at intersections were also shown to lead to incidental learning of spatial information in a virtual driving paradigm (Gramann et al., 2017).

Based on these results, we aimed at investigating the temporal scale of incidental spatial knowledge acquisition during assisted navigation. In two studies, we asked whether landmark-based instructions lead to long-term spatial learning of the navigated environment and how additional landmark information contribute to the incidentally acquired knowledge. This was tested by using standard and landmark-based auditory navigation instructions in a well-controlled simulator driving scenario (Gramann et al., 2017). The control group received standard navigation instructions as known from commercial navigation aids providing information about the distance to the upcoming intersection and the turning direction (e.g., “In 100 m, turn right at the intersection.”) Two versions of landmark-based navigation instructions were used in the first experiment including both the name of a landmark at the respective intersection and an additional information. In one instruction condition (personal-reference), this additional information included a reference to the participant’s personal interests. For example, the upcoming landmark was a bookstore, then the favorite author of the participant (e.g. J.R.R. Tolkien) was mentioned in the instructions: “Turn right at the bookstore. There, you can buy books of J.R.R. Tolkien”. All auditory navigation instructions were individualized for each participant in the personal-reference condition regarding different dimensions of personal interest ranging from food to cultural activities. In the other landmark-based instruction condition (contrast), the purpose of the additional information following the landmark reference was only to maintain the same length of the auditory instructions as provided in the personal-reference instruction condition. Thus, it contained redundant information, such as, “Turn right at the bookstore. There, you can buy books.” Auditory navigation instructions of all conditions were presented together with a semi-transparent arrow projected into the virtual environment indicating the upcoming turning direction. The instruction conditions used in the first experiment replicated the instructions used in the study by Gramann and colleagues (2017). In contrast to this earlier study, however, the present experiment added a three-week break between the assisted navigation task and the spatial tests.

In a second experiment, we then removed the visual route indicator to avoid shifts of attention towards this visual input that might lead to decreased spatial knowledge acquisition of the surrounding environment. In addition, another landmark-based instruction condition was used to test the impact of the information quality and quantity in the landmark-based instructions.

Based on previous studies, we expected a better spatial knowledge acquisition for navigators receiving landmark-based navigation instructions even when testing three weeks after navigating only one route through an unfamiliar environment.

## Experiment 1

Gramann and colleagues (2017) demonstrated an improved landmark and route knowledge when driving with landmark-based navigation instructions. To test whether this increase of incidentally acquired spatial knowledge was a short- or long-term effect and whether an advantage of the personal-reference instructions would appear only after a longer period of time, we replicated the experiment by adding a three-week delay in between assisted navigation and spatial tasks. The analysis of brain dynamics during cued-recall of landmark and route knowledge was described in a related paper (Wunderlich & Gramann, 2018).

### Methods

#### Participants

Forty-five participants (23 female) took part in the experiment. Gender distribution was balanced for all navigation instruction conditions. The age ranged from 19 to 37 years (M = 26.44 years, *SD* = 4.37 years). Participants were contacted directly or through an existing database and were reimbursed with 10 Euro per hour. All had normal or corrected to normal vision and owned a driver license for at least two years. They gave informed consent prior to experiment and the study was approved by the local research ethics committee.

#### Setup and Virtual Environment

The driving simulator was a Volkswagen Touran driver’s cabin with the virtual scene projected onto a white wall approximately two meters in front of the driver (see Figure 1a). Movements of the steering wheel and of the pedals were linked to the Game Controller, MOMO Racing Force Feedback Wheel (Logitech, Switzerland). This physical link restricted the maximum turning angle of the steering wheel to 110°. To enable sharp turns, the sensitivity of the controller was increased.

**Figure 1:**
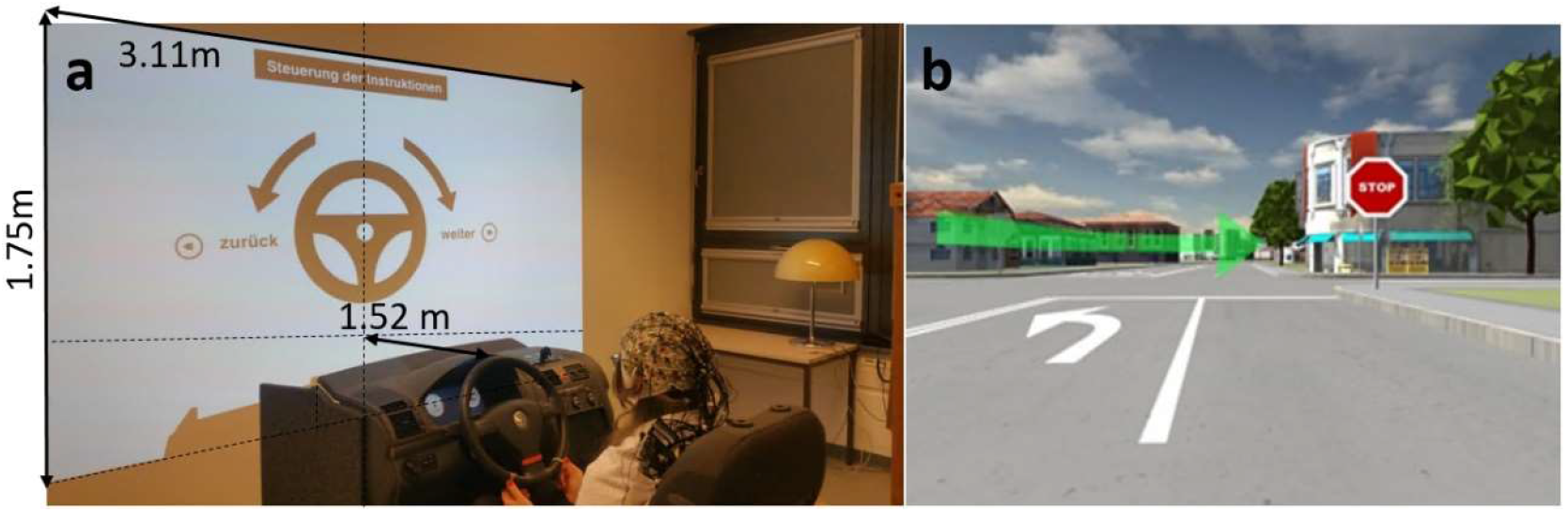
a) Driving simulator setup during task instructions and b) virtual scene during assisted navigation including the used hologram arrow projected into the virtual environment (Figure adapted from Wunderlich & Gramann, 2018).

We reused the virtual city model from Gramann and colleagues (2017) containing an unknown city covering 36 km^2^ with suburban and urban areas. In this simulation, traffic was reduced to one approaching car at the beginning of the route. The predefined route traversed once through the whole city including different speed limits and stop signs or traffic lights. At seven intersections, the route direction changed with an unpredictable order of turning direction. There were several buildings contrasting with all other buildings in the virtual environment. These landmarks were either located at *intersections* where the route direction changed and thus were referenced in the respective landmark-based navigation instruction or at intersections during *straight segments* of the route without accompanying navigation instructions.

About 100 m prior to a route direction change, the navigation instructions were initiated. Navigation instructions combined an auditory instruction with a visual cue. The latter was implemented by a semitransparent hologram arrow projected onto the environment comparable to a head-up display (see Figure 1b). To ensure that the participants remain on the predefined route, an automated resetting mechanism was in place which stopped the car after a wrong turn and positioned it back on the correct route facing the last intersection.

#### Study Design and Procedure

Participants were assigned to one out of three different navigation instruction conditions (standard, contrast, and personal-reference). The experiment was divided in two sessions separated by a three-week period (*M* = 21.07 days, *SD* = 1.65 days, *Min* = 20 days, *Max* = 24 days). During the first session, participants navigated the route assisted by the navigation instructions and in the second session, they solved a set of spatial tasks. Participants were not informed about the spatial tasks until the start of the second session to ensure incidental learning.

Prior to the assisted navigation session, all participants answered a pre-test questionnaire about their personal preferences sent to them via e-mail. For participants assigned to the personal-reference condition, the respective responses were included in the auditory navigation instructions using a text to speech software (Voice RSS, www.voicerss.org).

The assisted navigation session lasted one hour in total and started with the instructions about the traffic rules and a 5 min familiarization phase of self-determined driving through a very simple environment to accustom to the driving simulator setup. Thereafter, participants were further instructed to follow the navigation instructions when driving in the test environment. Following the 10 min ride, the participants’ task load was assessed using the National Aeronautics and Space Administration Task Load Index (NASA-TLX, Hart & Staveland, 1988). The affective state was assessed using the Affect Grid (Russell et al., 1989) and simulation sickness was recorded with three questions on a four-point Likert-scale (nausea, headache, and dizziness). Furthermore, participants answered several questionnaires targeting at driving and navigation habits, gaming experience, and spatial orientation (Santa Barbara Sense of Direction, SBSOD, Hegarty et al., 2002). At the end, the Reference Frame Proclivity Test (RFPT) was conducted to assess the individual preference for an egocentric or an allocentric reference frame during a path integration task (Goeke et al., 2015; Gramann et al., 2005).

The second experimental session after the 3-week break contained five tasks focusing on the incidentally acquired spatial knowledge during the assisted navigation. It started with a cued-recall task that contained snap-shots of 21 intersections in the environment taken from the drivers’ perspective and presented each one in a random order to the participant. In case the displayed landmark was located at an intersection with a direction change, participants were instructed to steer to the right or to the left according to the turning direction during assisted navigation. If the landmark was located at a straight route segment participants were instructed to push the gas pedal, and to apply the brakes in case of novel landmarks (landmarks that had not been experienced during the navigation phase but that were part of the VR environment). For each of the three landmark types seven landmarks were included. In the Scene Sorting task, participants were asked to remove the novel landmarks from the others and then sort the remaining ones according to their occurrence from start to end of the route. Afterwards, participants were handed an empty sheet of paper (DIN A3) and pens to draw a map of the route as well as everything they remembered from the environment. For the next task, participants had another familiarization phase in the driving simulator. Subsequently, they were placed into the test environment and instructed to drive the identical route as during the assisted navigation phase three weeks earlier but without any assistance. The automated resetting mechanism after incorrect turns ensured that participants reached the destination. Subsequently, the NASA-TLX, Affect Grid, and simulation sickness questionnaire were filled out. The final task of the test session was another sketch map drawing of the virtual environment to assess differences in spatial knowledge after being confronted with the environment for a second time. At the end of the session, participants provided feedback about the experiment and the landmark-based navigation instructions. Questionnaires throughout the test sessions, were presented and answered using a tablet (iPad 1, Apple Inc., Curtino, California) with the software LimeSurvey (Hamburg, Germany).

#### Statistics

Statistics were computed using the statistics software SPSS (International Business Machines Corporation (IBM) Analytics, Armonk, USA).

Group differences in individual measures were tested using one-way Analyses of Variance (ANOVAs) comparing the between-subject factor navigation instruction (standard, contrast, personal-reference).

Recognition sensitivity and number of free-recalled landmarks were tested in a mixed-measure ANOVA with the between-subject factor navigation instruction (standard, contrast, personal-reference) and the repeated measure factor landmark type (intersections, straight segments).

The percentage of correct direction responses to landmarks at intersections in the cued-recall task and the number of wrong turns during the second drive without navigation aid were tested in an univariate ANOVA comparing the levels of the between-subject factor navigation instruction condition (standard, contrast, personal-reference).

For significant main effects and interactions, post-hoc comparisons were computed using Bonferroni to correct for multiple comparisons. Partial eta squared was reported as an indicator for the effect size.

### Results

#### Individual Measures

Using several questionnaires and individual measures, we checked whether other factors might have impacted incidental spatial knowledge acquisition differentially for the two experimental groups.

The subjective mental load during assisted navigation was assessed with an univariate ANOVA and showed no differences navigation instruction conditions (*F*_(2,42)_ = 2.49, *p* = .095, 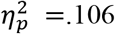; standard: *M* = 55.1, *SD* =20.4, contrast: *M* = 38.6, *SD* = 19.3, personal-reference: *M* = 45.4, *SD* = 21.2). This subjective measure was complemented by objective parameters of driving behavior. No group differences were revealed by testing the standard deviation of distance to the ideal line (vertical control) or the standard deviation in speed (horizontal control) with all *p*s > .445.

The subjective orienting ability was measured using the SBSOD resulting in a trend towards significance (*F*_(2,42)_ = 3.12, *p* = .055, 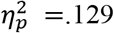). The means for the three instruction conditions showed slight variations with the standard instruction group showing the highest values (*M* = 4.01, *SD* = 0.87, contrast: *M* = 3.37, *SD* = 0.74, personal-reference: *M* = 3.83, *SD* = 0.55).

There were no group differences in the ratings of simulator sickness after the assisted navigation.

#### Landmark Knowledge

The cued-recall task primarily tested landmark knowledge. Presenting landmarks at intersections or straight segments as well as presenting novel landmarks allowed for the computation of the recognition sensitivity accounting for a potential individual response bias. The incorrect responses to novel landmarks served as a measure for the false alarm rate. Right and left turns of the steering wheel were both rated as hit in case of landmarks at intersections with a direction change. The mixed measures ANOVA testing recognition sensitivity with the between-subject factor navigation instruction condition and the repeated measures factor landmark type revealed a significant main effect of navigation instruction condition (*F*_(2,42)_ = 4.40, *p* = .018, 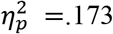) and a main effect of landmark type (*F*_(1,42)_ = 11.8, *p* = .001, 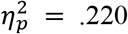). The interaction of both factors was also significant (*F*_(2,42)_ = 5.61, *p* = .007, 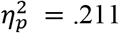). Bonferroni-corrected post-hoc comparisons of the interaction revealed that the recognition sensitivity for intersections in the standard instruction condition (*M* = -0.98, *SE* = 0.34) was significantly lower than in the landmark-based instruction conditions (contrast: *M* = 0.81, *SE* = 0.34, *p* = .002, personal-reference: *M* = 0.60, *SE* = 0.34, *p* = .006, all other *p*s > .849).

Another measure for landmark knowledge was the number of free-recalled landmarks in the second sketch map. This second sketch map was assessed at a comparable time point during the spatial test session as the sketch map of Gramann and colleagues (2017). For comparability reasons and because of the poor performance in the first sketch maps, we decided to present only the results of this second sketch map. The mixed-measure ANOVA for the number of correctly drawn landmarks in the second sketch map revealed a significant main effect of navigation instruction condition (*F*_(2,42)_ = 7.07, *p* = .002, 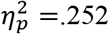). The main effect of landmark type (*F*_(1,42)_ = 86.0, *p* < .001, 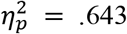) and the interaction of both factors were also significant (F(2,42) = 4.58, *p* = .025, 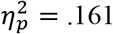). Post-hoc comparisons revealed less recalled landmarks at intersections in the standard instruction condition (*M* = 2.87, *SE* = 0.40) compared to the landmark-based instruction conditions (contrast: *M* = 5.27, *SE* = 0.40, *p* < .001; personal-reference: *M* = 4.88, *SE* = 0.40, *p* = .003). Free-recall of landmarks at straight segments was comparable across conditions *p*s > .116.

#### Route Knowledge

The cued-recall task measures landmark as well as route knowledge. Because the responses to landmarks also required navigation decisions, this test can also assess route knowledge (or Heading Orientation according to Huang et al., 2012) including stimulus-response associations. The univariate ANOVA testing the percentage of correct responses to landmarks at intersections showed a main effect of instruction condition (*F*_(2,42)_ = 11 .1, *p* < .001, 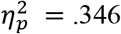). Participants in the control group performed worse (*M* = 25.7%, *SE* = 3.68%) compared to both landmark-based navigation instruction conditions (contrast: *M* = 44.8%, *SE* = 3.68%, *p* = .002, personal-reference: *M* = 48.6%, *SE* = 3.68%, *p* < .001). The two landmark-based instruction condition led to comparable route knowledge (*p* = .367).

The number of wrong turns during driving the route without navigation assistance was tested in an univariate ANOVA and revealed comparable numbers of incorrect turns across all three navigation instruction conditions (*F*_(2,42)_ = 1.64, *p* = .207, 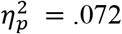).

### Discussion

The first experiment replicated the setup and paradigm of Gramann and colleagues (2017) delaying the spatial task session by three weeks. The results revealed better landmark and route knowledge for landmark-based navigation instructions even after three weeks with no significant advantage of navigation instruction including references to personal interest as compared to contrast landmark-based instructions.

The cued-recall task was a combination of landmark recognition and route direction recall. For landmarks alongside the route, it required retrieval of route information besides a simple recognition of the landmark. Consequently, after successfully recognizing a landmark at an intersection with route direction change, there were still two possible response options (right and left). In contrast, landmarks at straight segments and novel landmarks had only one response option. This was considered in the analysis of recognition sensitivity by joining both turning responses as hit for landmarks at intersections. In contrast, the analysis of route knowledge considered only the correct direction as correct response.

When checking for potential differences in individual measures between the groups, the SBSOD and the subjective mental load revealed trends with p-Values below . 10. The mental load subscale of the NASA-TLX had been used as covariate in the analyses of Gramann and colleagues (2017). However, in none of their analyses of covariance (ANCOVAs) a related main effect or interaction reached significance. For the reported dataset, we further tested for significant correlations of the NASA-TLX and the SBSOD with all dependent variables and included the measure as covariate in the respective analyses. Because no significant effects of the two covariates were revealed, we reported the results without covariates to be consistent in both experiments. Based on the absence of any influence of individual mental load or sense of direction in the analyses on group-level, the changes in incidental spatial knowledge acquisition have to be due to variations in the navigation instructions.

The first experiment demonstrated improved spatial knowledge acquisition when participants navigated through an unfamiliar environment using landmark-based navigation instructions. This incidental learning of landmarks and route information was observed even three weeks after a single encounter of an unfamiliar environment, replicating previous results that demonstrated an immediate spatial knowledge acquisition (Gramann et al., 2017). However, the standard instructions directed participants’ attention towards the “intersection” that was further overlaid by the visual navigation cue projected into the virtual environment. Possibly, this combination of auditory instruction combined with a visual cue in the central field of view was more detrimental for the standard navigation instruction condition lacking an environmental cue that could draw attention to areas around the visual cue. Thus, in experiment 2, the visual route indicator was removed to rule out any potential inhibitory impact on spatial knowledge acquisition.

## Experiment 2

To exclude any impact of additional visual instructions on spatial knowledge acquisition during navigation, the second experiment tested whether the overlay of the previously used visual navigation cue might have caused the observed reduction of landmark knowledge at intersections. Previous studies investigating the impact of landmark-based navigation instructions on spatial knowledge acquisition used auditory navigation instructions alongside a visual navigation cue - a semi-transparent hologram arrow projected in the virtual environment directing the route direction (see Figure 1b; Gramann et al., 2017; Wunderlich & Gramann, 2018).

Furthermore, the previously used landmark-based navigation instructions did not allow for further dissociating the impact of the additional information in the navigation instructions on spatial knowledge acquisition. One explanation for improved spatial knowledge acquisition compared to standard instructions could have been the personal-reference in the instruction. Alternatively, simply naming a landmark and adding more detailed information might have led to the observed improved spatial knowledge. To overcome this limitation, the second experiment introduced a new landmark-based navigation instruction condition, the *long* instruction condition. Long landmark-based navigation instructions referenced a landmark and provided additional detailed information about this landmark. This additional information contained more semantic information than the redundant description of the landmark in the contrast navigation instruction, but did not relate to the personal interests of the participants. Thus, there was no need to individualize the additional information and all participants of the long condition heard the same navigation instructions (e.g. “Turn right at the bookstore. There, public readings take place every week.”).

With the second experiment, we aimed to replicate the positive effect of landmark-based navigation instructions on incidental spatial knowledge acquisition. Changes in the experimental protocol targeted the impact of the visual cue as well as the impact of the content of the additional information included in the landmark-based instructions.

### Methods

#### Participants

The second experiment comprised 29 participants (13 females) with gender balanced across experimental conditions (standard navigation instruction with 16 participants, 7 female; long navigation instruction with 13 participants, 6 female). The age ranged between 22 and 53 years (*M* = 31.9 years, *SD* = 6.40 years). Participants were recruited through an existing database or personal contact and were reimbursed (8 Euro per hour) or received course credit. Participants were required to have had a drivers’ license for two years or more and to have driven at least 1000 km per year to assure a basic driving experience. All had normal or corrected to normal vision and gave informed consent prior to the study. The study was approved by the local ethics committee.

#### Measurement and Apparatus

Only slight changes were made regarding the technical setup of the driving simulator used in experiment 1. Driving experience was improved by allowing for turning the steering wheel 360 degrees from neutral position in both directions.

The identical virtual city and route was used. In contrast to the previous studies, the visual turning indicator (Figure 1b) was replaced by a second auditory turning instruction (e.g. “Now turn right.”) which was played when reaching the respective intersection. The second navigation instruction was presented at all intersections with a route direction change and was identical for all navigation instruction conditions.

#### Study Design and Procedure

The period between the navigation session and the spatial tests was again three weeks. Two groups of participants received either standard or long landmark-based navigation instructions. Compared to the spatial task order in experiment 1, the first sketch map drawing was moved to be the first spatial test followed by the other tasks accordingly. Changing the order of spatial tasks allowed for unbiased free recall measures as they might have been influenced by proceeding tasks in experiment 1. All other task characteristics were kept constant, only some questionnaires were added to or removed from the paradigm.

#### Statistics

Statistics of the second experiment were analog to the first experiment with only two levels of the between-subject factor navigation instruction condition (standard, long).

### Results

#### Individual Measures

The subjective mental load did not vary significantly between navigation instruction conditions when tested in an univariate ANOVA (*F*_(1,27)_ = 2.62, *p* = .117, 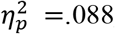, standard: *M* = 41 .9, *SD* = 23.7, long: *M* = 55.9 .4, *SD* = 22.4). The respective analyses of the driving parameters representing vertical and horizontal control were also not significant (*p*s > .308).

Subjective orienting ability as assessed with the SBSOD was comparable for all experimental groups (*F*_(1,27)_ = 1.13, *p* = .297, 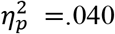) with *M* = 3.61, *SD* = 0.84 (standard: *M* = 3.76, *SD* = 0.90, long: *M* = 3.43 *SD* = 0.75).

There was no group difference for the simulator sickness ratings.

#### Landmark Knowledge

The mixed measures ANOVA of recognition sensitivity with the between-subject factor navigation instruction condition (standard, long) and the repeated measures factor landmark type (intersections, straight segments) revealed a significant main effect of navigation instruction condition (*F*_(1,27)_ = 8.14, *p* = .008, 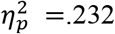) and a main effect of landmark type (*F*_(1,27)_ = 7.19, *p* = .012, 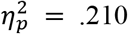). The interaction of both factors also reached significance (*F*_(1,27)_ = 5.39, *p* = .009, 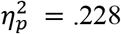). Bonferroni-corrected post-hoc comparisons of the interaction revealed a significant difference in recognition sensitivity between the control group (*M* = -0.40, *SE* = 0.31) and landmark-based instructions (*M* = 1.24, *SE* = 0.34, *p* = .001) for landmarks at intersections with direction changes, whereas recognition sensitivity for landmarks at straight segments was comparable (*p* = .276).

In the analysis of the number of correctly drawn landmarks in the second sketch map, the main effect of navigation instruction condition did not reach significance (*F*_(1,27)_ = 3.44, *p* = .075, 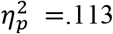). The main effect of landmark type (*F*_(1,27)_ = 34.8, *p* < .001, 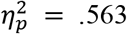) and the interaction of both factors were significant (*F*_(1,27)_ = 5.85, *p* = .023, 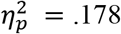). Post-hoc comparisons revealed less recalled landmarks at intersections in the standard instruction condition (*M* = 2.81, *SE* = 0.43) compared to the long landmark-based instruction condition (*M* = 4.46, *SE* = 0.48, *p* = .016). Free-recall of landmarks at straight segments was comparable across conditions *p* = .736.

#### Route Knowledge

The univariate ANOVA of percentage correct responses to landmarks at intersections comparing the between-subject factor navigation instruction condition (standard, long) was significant (*F*_(1,27)_ = 11.8, *p* = .002, 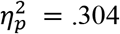). Participants in the control group performed worse (*M* = 29.5%, *SE* = 3.90%) than the landmark-based navigation instruction condition (*M* = 49.5%, *SE* = 4.33%).

The number of wrong turns during driving the route without navigation assistance was tested in an univariate ANOVA comparing the navigation instruction conditions (F(1,27) = 9.30, *p* = .005, 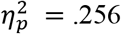). Participants who had previously navigated with standard instructions turned more often into a wrong direction (*M* = 7.19, *SE* = 0.51) compared to those who had used long landmark-based instructions (*M* = 4.85, *SE* = 0.57).

### Discussion

Experiment 2 aimed at investigating whether the previously used visual turn indicator partially overlapping with the environment prevented incidental spatial knowledge acquisition especially in combination with the standard navigation instructions. To this end, the former setup was replicated while replacing the visual navigation instruction with a second, short auditory turn instruction. Furthermore, we introduced the long landmark-based navigation instruction combining the landmark name and additional detailed information about the respective landmark.

Group means of individual measures did not differ between navigation instruction conditions allowing for the assumption that differences in the spatial knowledge were associated with the respective auditory navigation instructions. The results of experiment 2 showed a better landmark and route knowledge for the long navigation instruction condition compared to standard instructions replicating the positive impact of landmark-based, auditory navigation instructions.

A lower recognition sensitivity was observed in the standard navigation instruction condition when responding to landmarks located at intersections with route direction changes. This effect was observed even after removing the visual cue replicating the results of experiment 1 and the earlier study (Gramann et al., 2017) and confirming the hypothesis that landmark-based navigation instructions foster incidental spatial knowledge acquisition. It can be concluded that the visual turn indicator was not the cause for the previously observed reduced landmark knowledge in the standard navigation instruction condition. In the present study, no visual navigation instruction was presented that might have prevented information processing of environmental features at intersections with route direction changes. Still, navigators receiving standard navigation instructions demonstrated less landmark and route knowledge acquisition. The differences in landmark knowledge must therefore be related to differences in the auditory navigation instructions. The standard navigation instructions referring to the next intersection allocate the navigators’ attention towards the street whereas the landmark-based navigation instructions draw attention to specific environmental features surrounding the intersection and thus maintain information processing of environmental aspects.

As there were only minor changes in the general setup, a direct comparison of the performance results with the data of experiment 1 is possible. Both experiments had a break of three weeks between the navigation phase and the spatial tests. The performance of both standard navigation instruction conditions revealed comparable results for all dependent measures indicating the reliability of the navigation instruction effects. In addition, participants receiving long landmark-based navigation instructions revealed a comparable performance as participants using landmark-based navigation instructions with references to personal interests. The comparable performance for long and personal-reference instructions supports the assumption that improved spatial knowledge acquisition through landmark-based navigation instructions was based on additional semantic information related to navigation relevant landmarks and not to the personal reference. Simply mentioning a personal interest might not be sufficient to trigger the personal-reference effect (for a review see Symons & Johnson, 1997). Beneficially, this reduces the requirement to collect personal information from the user and related concerns regarding data security. Furthermore, as improved incidental spatial knowledge acquisition was based on the landmark-based auditory navigation instructions and not related to the visual navigation indicator, the applicability of this simple, but promising modification of navigation assistance systems should be further tested in different navigation contexts.

## General Discussion

Two experiments replicated and slightly adapted the setup and paradigm of Gramann and colleagues (2017). This allowed to address open questions resulting from their initial findings. In the first experiment, we investigated the long-term impact of incidental spatial knowledge acquisition based on landmark-based navigation instructions. In the second experiment, the impact of the visual cues and the content of landmark-based instructions was addressed.

The results of both experiments replicated the positive impact of augmenting landmarks during assisted navigation on spatial knowledge acquisition (Goodman et al., 2005; Gramann et al., 2017; Li et al., 2014; Löwen et al., 2019). An increased landmark and route knowledge for landmark-based navigation instructions was consistently revealed despite all setup changes and individual differences.

Even after a break of three weeks, the positive impact of landmark-based instructions on incidentally acquired landmark and route knowledge was replicated in both experiments. When comparing the number of incorrect turning decisions during the second drive, the significant difference of landmark-based versus standard instructions as shown by Gramann and colleagues (2017) was only replicated in the second experiment. However, the direction of the effect was identical in both experiments without reaching significance in experiment 1.

The results further provided first evidence that long landmark-based instructions lead to a comparable level of landmark learning as landmark-based instructions that include personal-references. This finding allows to standardize navigation instructions for different users without requiring individualized modifications. The long landmark-based instructions do not necessitate the use of personal information which would otherwise be associated with data security concerns. The results from the reported experiment demonstrate that it would be sufficient to access publicly available information and then generate the landmark-based navigation instructions. This was already done for the inclusion of landmark names in navigation instruction by other researchers (Dräger & Koller, 2012; Rousell & Zipf, 2017).

Furthermore, a previous limitation of the test setup was overcome. Based on the results we can conclude that the visual instructions did not cause the reduced spatial knowledge acquisition at intersections with route direction changes.

A remaining limitation of the findings is that none of the tasks allowed conclusions about incidentally acquired survey knowledge. Even though sketch map drawings were included, the quality of the drawings was too low for the analysis of angular and distance measures. Navigating only once through an unfamiliar environment seems to be insufficient to reproduce a map-like representation of an environment.

## Conclusions

Both reported experiments replicate the previously described positive impact of landmark-based navigation instructions on incidental acquisition of landmark and route knowledge. Results revealed that the effect endures over longer time periods and ruled out that it was induced by the former provided visual cue. Thus, landmark-based navigation assistance systems help to maintain the processing and extraction of navigation relevant information from the environment.

Future research should test this effect when using landmark-based navigation aids multiple times enabling the investigation of incidentally acquired survey knowledge. Furthermore, the setup should be transferred to more realistic settings and other locomotion modes to test the ecological validity. Additionally, further investigation of accompanied gaze and brain activity would allow deeper understanding of the underlying physiological changes and involved cognitive processes.

## Acknowledgements

This research was supported by a stipend from the Stiftung der Deutschen Wirtschaft to AW. We would like to thank Prof. Matthias Rötting at TU Berlin for providing the driving simulator setup. Additionally, we give thanks to Christopher Hahn and Juliane Kayser for helping to prepare and conduct the second experiment.

The authors declare that there is no conflict of interest regarding the publication of this paper.

Data is available on request from the corresponding author.

## References

Aporta, C., & Higgs, E. (2005). Global positioning systems, inuit wayfinding, and the need for a new account of technology. Current Anthropology, 46(5), 729–753. https://doi.org/10.1086/432651

Baus, J., Kray, C., & Krüger, A. (2001). Visualisation of route descriptions in a resource-adaptive navigation aid. Cognitive Processing, 2, 323–345.

Dräger, M., & Koller, A. (2012). Generation of landmark-based navigation instructions from open-source data. EACL 2012 - 13th Conference of the European Chapter of the Association for Computational Linguistics, Proceedings, 757–766.

Evans, G. W., Smith, C., & Pezdek, K. (1982). Cognitive maps and urban form. Journal of the American Planning Association, 48(2), 232–244. https://doi.org/10.1080/01944368208976543

Fenech, E. P., Drews, F. A., & Bakdash, J. Z. (2010). The effects of acoustic turn-by-turn navigation on wayfinding. Proceedings of the Human Factors and Ergonomics Society, 3, 1926–1930. https://doi.org/10.1518/107118110X12829370263287

Gardony, A. L., Brunyé, T. T., Mahoney, C. R., & Taylor, H. A. (2013). How Navigational Aids Impair Spatial Memory: Evidence for Divided Attention. Spatial Cognition and Computation, 13(4), 319–350. https://doi.org/10.1080/13875868.2013.792821

Gardony, A. L., Brunyé, T. T., & Taylor, H. A. (2015). Navigational Aids and Spatial Memory Impairment: The Role of Divided Attention. Spatial Cognition and Computation, 15(4), 246–284. https://doi.org/10.1080/13875868.2015.1059432

Goeke, C., Kornpetpanee, S., Köster, M., Fernández-Revelles, A. B., Gramann, K., & König, P. (2015). Cultural background shapes spatial reference frame proclivity. Scientific Reports, 5, 1—13. https://doi.org/10.1038/srep11426

Goodman, J., Brewster, S. A., & Gray, P. (2005). How can we best use landmarks to support older people in navigation? Behaviour and Information Technology, 24(1),3–20. https://doi.org/10.1080/01449290512331319021

Gramann, K., Hoepner, P., & Karrer-Gauss, K. (2017). Modified Navigation Instructions for Spatial Navigation Assistance Systems Lead to Incidental Spatial Learning. Frontiers in Psychology, 8(FEB). https://doi.org/10.3389/fpsyg.2017.00193

Gramann, K., Müller, H. J., Eick, E. M., & Schönebeck, B. (2005). Evidence of separable spatial representations in a virtual navigation task. Journal of Experimental Psychology: Human Perception and Performance, 31(6), 1199–l1223. https://doi.org/10.1037/0096-1523.31.6.1199

Hart, S. G., & Staveland, L. E. (1988). Development of NASA-TLX (Task Load Index): Results of Empirical and Theoretical Research. Advances in Psychology, 52(C), 139–183. https://doi.org/10.1016/S0166-4115(08)62386-9

Hegarty, M., Richardson, A. E., Montello, D. R., Lovelace, K., & Subbiah, I. (2002). Development of a self-report measure of environmental spatial ability. Intelligence, 30(5), 425–447. https://doi.org/10.1016/S0160-2896(02)00116-2

Huang, H., Schmidt, M., & Gartner, G. (2012). Spatial knowledge acquisition with mobile maps, augmented reality and voice in the context of GPS-based pedestrian navigation: Results from a field test. Cartography and Geographic Information Science, 39(2), 107–116. https://doi.org/10.1559/15230406392107

Klippel, A., Hirtle, S., & Davies, C. (2010). You-Are-Here Maps: Creating Spatial Awareness through Map-like Representations. Spatial Cognition & Computation, 10(2–3), 83–93. https://doi.org/10.1080/13875861003770625

Li, R., Fuest, S., & Schwering, A. (2014). The effects of different verbal route instructions on spatial orientation. The 17th AGILE Conference on Geographic Information Science, 3–6.

Löwen, H., Krukar, J., & Schwering, A. (2019). Spatial learning with orientation maps: The influence of different environmental features on spatial knowledge acquisition. ISPRS International Journal of Geo-Information, 8(3). https://doi.org/10.3390/ijgi8030149

Mosier, K. L., Skitka, L. J., Burdick, M. D., & Heers, S. T. (1996). Automation Bias, Accountability, and Verification Behaviors. Proceedings of the Human Factors and Ergonomics Society Annual Meeting, 40(4), 204–208. https://doi.org/10.1177/154193129604000413

Parasuraman, R. (2000). Designing automation for human use: Empirical studies and quantitative models. Ergonomics, 43(7), 931–951. https://doi.org/10.1080/001401300409125

Parush, A., Ahuvia, S., & Erev, I. (2007). Degradation in spatial knowledge acquisition when using automatic navigation systems. Lecture Notes in Computer Science (Including Subseries Lecture Notes in Artificial Intelligence and Lecture Notes in Bioinformatics), 4736 LNCS, 238–254. https://doi.org/10.1007/978-3-540-74788-8_15

Rousell, A., & Zipf, A. (2017). Towards a landmark-based pedestrian navigation service using OSM data. ISPRS International Journal of Geo-Information, 6(3). https://doi.org/10.3390/ijgi6030064

Russell, J. A., Weiss, A., & Mendelsohn, G. A. (1989). Affect Grid: A single-item scale of pleasure and arousal. Journal of Personality and Social Psychology, 57(3), 493–502. https://doi.org/10.1037/0022-3514.57.3.493

Symons, C. S., & Johnson, B. T. (1997). The Self-Reference Effect in Memory: A Meta-Analysis. Psychological Bulletin, 121(3), 371–394. https://doi.org/10.1037/0033-2909.121.3.371

Wunderlich, A., & Gramann, K. (2018). Electrocortical evidence for long-term incidental spatial learning through modified navigation instructions. In Lecture Notes in Computer Science (including subseries Lecture Notes in Artificial Intelligence and Lecture Notes in Bioinformatics): Vol. 11034 LNAI. Springer International Publishing. https://doi.org/10.1007/978-3-319-96385-3_18

